# Predicting Synthetic Lethal Interactions using Heterogeneous Data Sources

**DOI:** 10.1101/660092

**Authors:** Herty Liany, Anand Jeyasekharan, Vaibhav Rajan

## Abstract

**Motivation:** A synthetic lethal (SL) interaction is a relationship between two functional entities where the loss of either one of the entities is viable but the loss of both entities is lethal to the cell. Such pairs can be used as drug targets in targeted anticancer therapies, and so, many methods have been developed to identify potential candidate SL pairs. However, these methods use only a subset of available data from multiple platforms, at genomic, epigenomic and transcriptomic levels; and hence are limited in their ability to learn from complex associations in heterogeneous data sources.

**Results:** In this paper we develop techniques that can seamlessly integrate multiple heterogeneous data sources to predict SL interactions. Our approach obtains latent representations by collective matrix factorization based techniques, which in turn are used for prediction through matrix completion. Our experiments, on a variety of biological datasets, illustrate the efficacy and versatility of our approach, that outperforms state-of-the-art methods for predicting SL interactions and can be used with heterogeneous data sources with minimal feature engineering.

**Availability:** Software available at https://github.com/lianyh

## 1 Introduction

Genomic studies have shed light on several aspects of cancer, from the understanding of how the disease initiates and progresses to genomic drivers of the disease and the development of first generation of targeted therapies (Hyman *et al.*, 2017). Cancer develops as a result of mutational events caused by endogenous and exogenous process; these mutations enable cancer cells to gain selective advantage over healthy cells resulting in uncontrolled proliferation and ultimately metastasis (Hanahan and Weinberg, 2011). Large-scale molecular profiling of major cancer types have been completed (Hudson *et al.*, 2010). Multi-omics data, including copy number, gene expression, DNA methylation, microRNA and clinical data of several cancers have been collected and analyzed, for example in the Cancer Genome Atlas Research Network (Weinstein *et al.*, 2013). A key challenge of cancer studies is in the integration of data generated on different platforms and at different levels – genomic, epigenomic and transcriptomic levels (Senft *et al.*, 2017).

Extensive studies of the genomic landscape of tumors have revealed vulnerabilities that have been fruitfully exploited to develop *targeted therapeutics* that offer highly specific therapies with fewer adverse effects and the potential to reduce overtreatment (O’Neil *et al.*, 2017). One promising direction has been the use of *synthetic lethality* for developing drug targets. A **synthetic lethal (SL)** genetic interaction is a functional relationship between two genes or functional entities where the loss of either entity is viable but the loss of both is lethal. SL pairs have been exploited in targeted cancer therapeutics: the basic idea is that in a malignant cell, functionally disruptive mutation in one of the two genes (say, A) of an SL pair (A,B) leads to dependency on B for survival and cancer cells can be selectively killed by inhibiting B. Non-cancerous cells, that have A, survive even when B is inhibited. See fig. 1 for a schematic. For example, mutations causing functional loss of BRCA1/2 genes leads to deficiency of DNA Damage Response mechanism and dependence on the protein PARP1/2 (Bryant *et al.*, 2005). Drugs based on PARP inhibitors are found to be promising in the treatment of breast cancer (Tutt *et al.*, 2009) and ovarian cancer (Audeh *et al.*, 2010). However, such SL interactions in humans remain largely unknown and there is a need for new methods to discover such pairs.

**Fig. 1:**
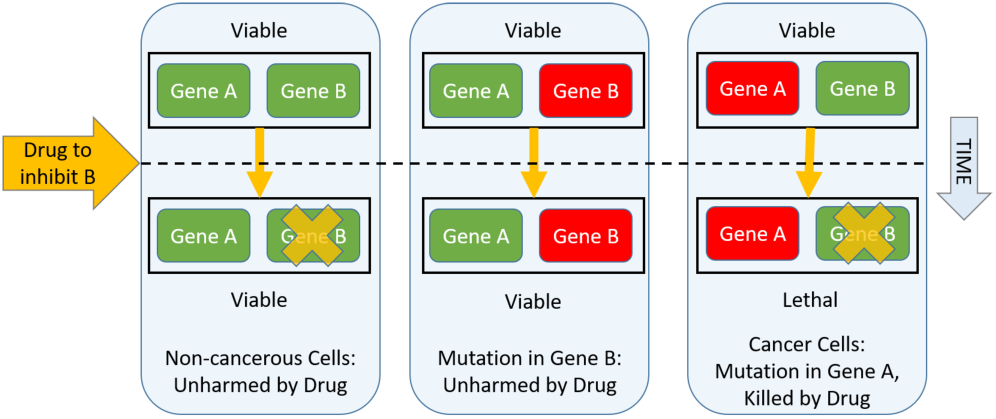
Synthetic Lethality: Genes A and B enable functionally redundant mechanisms and any one of them can ensure cell survival. If such pairs are found in cancer with one of the genes mutated, then the other can be targeted for developing highly specific drugs.

Synthetic Lethality has been considered a foundation for development of targeted anticancer therapies (Brough *et al.*, 2011; Senft *et al.*, 2017). As a result, large number of screens have been developed, such as RNA interference screens and CRISPR screens to identify potential SL pairs. Although such screens are effective approaches, they are costly and labour-intensive and significant challenges remain: first, since these genetic interactions are lethal, mutant recovery and identification become difficult; second, many SL pairs are conditionally dependent and may not be conserved in all genetic backgrounds or in different cellular conditions and third, large number of SL pairs need to be queried to identify SL interactions (O’Neil *et al.*, 2017). These genome-wide screens typically scan a few thousand candidate pairs of just one ‘anchor’ cancer driver gene of interest; due to the large combinatorial space of pairwise interactions, only a small fraction have been analyzed (Senft *et al.*, 2017).

Computational methods have been developed to identify potential SL pairs, reducing the number of candidates that can be functionally analyzed through genome-wide screens. These include machine learning based methods to predict genetic interactions in different species (Costanzo *et al.*, 2010; Lu *et al.*, 2013), in cancer (using yeast SL pairs) (Conde-Pueyo *et al.*, 2009; Srivas *et al.*, 2016), using metabolic modeling (Folger *et al.*, 2011; Frezza *et al.*, 2011), using evolutionary characteristics (Lu *et al.*, 2013; Srivas *et al.*, 2016), using transcriptomic profiles (Kim *et al.*, 2016) and by mining cancer patient data (Jerby-Arnon *et al.*, 2014; Sinha *et al.*, 2017; Lee *et al.*, 2018). All of these methods use only a subset of available data from multiple platforms, at genomic, epigenomic and transcriptomic levels. Individual analysis of the data sources may not reveal critical associations and potential causal relationships and there is a need to develop drug target discovery methods that can effectively integrate the diverse data sources that describe cancer at different levels.

Most biological datasets can be represented as matrices, where a matrix contains pairwise relational data between two *entities*. For example, a gene expression data matrix represents a relationship between entities, patients and genes. Thus, a collection of matrices may have multiple relationships between entities and each entity may be involved in multiple relationships. Collective Matrix Factorization (CMF) (Singh and Gordon, 2008), and extensions thereof, are models designed to collectively learn from multiple such relationships. These models generalize the idea of matrix factorization to a collection of matrices. They learn a latent representation for each entity in a way that information from multiple matrices are integrated seamlessly. These entity-specific latent representations can then be used in predictive tasks. However, CMF cannot model collections of matrices where there are two or more matrices describing the relation between the same entity, e.g., pairwise gene co-expression and mutual exclusivity information, that both contain relationships between the same entity, genes. Many biological datasets represent such relations, including the relation of synthetic lethality.

In this paper, we develop techniques to model arbitrary collections of matrices, that include two or more relations between the same entity. This extends the modeling capability of CMF to a much larger set of heterogeneous biological data. We evaluate our techniques in the task of predicting synthetic lethality for a pair of genes. We compare our techniques with four different collections of data, used by state-of-the-art methods for SL prediction. These methods involve the development of task-specific statistical inference tests or sophisticated feature engineering. Our CMF-based techniques can be used with derived features as well as the input data directly, with minimal feature engineering, and in each of the four cases, our techniques match or outperform previous methods, thereby demonstrating the accuracy and versatility of our method.

## 2 Related work

A comprehensive review of methods based on machine learning and network interaction can be found in (Madhukar *et al.*, 2015). In this section we provide a brief overview of some recent statistical and machine learning based approaches for predicting SL pairs.

### Statistical Approaches

*DAISY* applies three statistical inference procedures to identify potential SL pairs (Jerby-Arnon *et al.*, 2014). The first strategy, called genomic survival of the fittest (SoF), uses Somatic Copy Number Alteration (SCNA) and gene expression data to detect significantly infrequent co-inactivations in gene pairs. The second strategy, uses shRNA essentiality screens, SCNA and gene expression profiles, to identify pairs where inactivity or over-activity of a gene induces essentiality of the partner gene. The third test checks for significant pairwise co-expression in transcriptomic data, since SL pairs, participating in related biological processes are likely to be coexpressed. A gene pair is considered SL if all three criteria are satisfied.

In a similar approach, *ISLE* uses lab-screened SL pairs as inputs and identifies those pairs that are predictive of patients’ drug response (Lee *et al.*, 2018). Thus, ISLE can be viewed as a filtering algorithm to obtain clinically relevant SL pairs, from an initial (larger) collection of potential SL pairs. They apply three statistical procedures. In the first procedure, gene expression and SCNA data is used to identify candidate gene pairs with significantly infrequent co-inactivations. Second, a gene pair is selected if its co-inactivation leads to better predicted patient survival compared to when it is not co-inactivated. Survival probability is predicted using Cox proportional hazard model. Third, pairs with high phylogenetic similarity are identified, since functionally interacting genes tend to co-evolve. The final output consists of those pairs that fulfill all three criteria. Thus, apart from SCNA, gene expression and gene essentiality profiles, ISLE also uses clinical data and phylogeny information.

### Machine Learning Approaches

Ensemble-based classifiers have been used in many models to predict SL pairs, using both yeast and human data. For example, Pandey *et al.* (2010) developed the Multi-Network and Multi-Classifier (*MNMC*) framework to predict SL interactions in yeast, using six classifiers and several features extracted from PPI networks, transcription factor bindings, functional annotations, mutant phenotype data, phylogenetic profiles of proteins, KEGG pathway memberships of genes, sequence similarity and gene network modules and clique communities. *MetaSL*, developed by Wu *et al.* (2014), also used an ensemble of classifiers, that learnt the relative weight of each classifier in the ensemble and was found to outperform MNMC in predicting yeast SL pairs. They extracted features for their classifier from PPI networks, gene ontologies, gene expression and various similarity scores based on co-complex membership, co-pathway membership, whether or not they are paralogs, the number of their common or interacting domains as well as affinity in mass-spectrometry purifications. They did not predict human SL pairs directly, but through orthologous mapping from yeast to human genes. In a study that directly predicted human SL pairs, Lu *et al.* (2015) also use an ensemble of multiple classifiers (that we call *MCA*) using five features derived from Copy Number Variation (CNV) and RNASeq data. The features measure homozygous, heterozygous and mixed co-loss events as well as co-under-expression and co-inverse-expression events.

Note the heterogeneity of data sources used in all these methods. Further, the design of each of these methods depends on the data used. For example, the statistical tests chosen in DAISY or ISLE may have to be modified if different or additional data sources are used. Considerable effort has been devoted to designing relevant features in the machine learning methods, where feature engineering depends on the data used. None of these methods can seamlessly integrate arbitrary collections of heterogeneous data sources for predicting SL pairs.

## 3 Background

In this section we briefly describe CMF and its limitation with respect to modeling heterogeneous biological datasets.

For a single matrix *X* ∈ ℝ^*p*×*q*^, a low-rank factorization aims to obtain latent factors *U* ^(1)^ ∈ ℝ^*p*×*K*^, *U* ^(2)^ ∈ ℝ^*q*×*K*^, such that 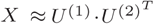, where the *K* < min(*p, q*). The factors are learnt by solving the optimization problem: 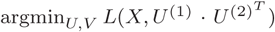, where *L* denotes a loss function.

Collective Matrix Factorization (CMF), proposed by Singh and Gordon (2008), generalizes the idea of factorization to an arbitrary collection of matrices. CMF aims to jointly obtain low-rank factorizations of arbitrary collections of *M* matrices (indexed by *m*), 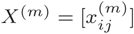, that describe relationships between *E* entities (*e*_1_, … *e*_*E*_), each with dimension 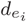. The entities corresponding to the rows and columns of the *m*^*th*^ matrix are denoted by *r*_*m*_ and *c*_*m*_ respectively. Each matrix is approximated by product of low rank-*K* factors that form the representations of the associated row and column entities: 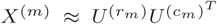 where 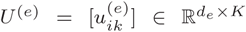 is the low-rank matrix for entity type *e*. Any two matrices sharing the same entity use the same low-rank representations as part of the approximation, which enables sharing information. A link function *f* may be applied to model non-linear relationships: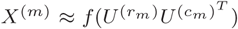. The latent factors are learnt by solving the optimization problem:

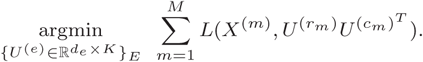

A regularizer is additionally used in some formulations. Solutions to this optimization problem obtained through Stochastic Gradient Descent have been found to yield good performance (Bouchard *et al.*, 2013).

Consider the example shown in fig. 2. The matrices could represent clinical data (*X*_1_), gene expression data (*X*_2_), and phylogenetic profiles (*X*_3_). Each matrix describes the relation between two entities, along its two dimensions. The entities in this example are patients (*e*_1_), clinical features (*e*_2_), genes (*e*_3_), and species (*e*_4_). CMF can learn entity-specific latent factors 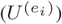 which are learnt collectively from all three matrices (with 4 entities): 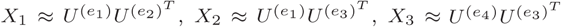. Due to this formulation, latent representations (e.g., 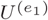) of entities that are shared across matrices (*e*_1_ across *X*_1_, *X*_2_) are learnt from all the matrices containing that entity and indirectly from other entities. Note that CMF can be used to learn entity-specific latent representations from *any* number of input matrices.

**Fig. 2:**
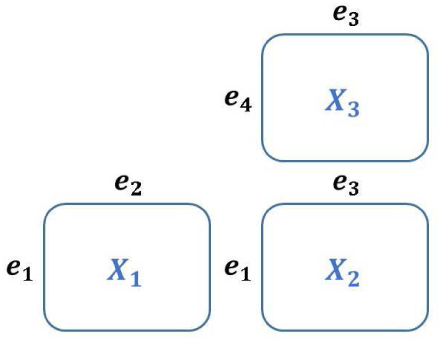
Example with 3 matrices (*X*_1_, *X*_2_, *X*_3_) and 4 entities (*e*_1_, *e*_2_, *e*_3_, *e*_4_). CMF (and gCMF) learns latent factors for each entity by collectively using all the information in any collection of matrices.

Group-Sparse CMF (gCMF) extends the CMF formulation through the use of group-sparse priors (Klami *et al.*, 2014). Individual matrices may have structured noise independent of other matrices, that cannot be captured by the element-wise noise terms. To model such noise, automatic creation of private factors is enabled by placing group-sparse priors on the columns of the matrices of *U* ^(*e*)^. If the *k*^*th*^ column of *U* ^(*e*)^ is null for all but two entity types *r*_*m*_ and *c*_*m*_, then the *k*^*th*^ factor is private to relation *m* since it impacts only matrix *X*^(*m*)^. If more than two groups of variables are non-zero then the factor is private to a subset of entities. The complete probabilistic model and Variational Bayesian inference for both Gaussian and non-Gaussian observations are presented in (Klami *et al.*, 2014).

CMF is an unsupervised learning method, but it can be used for matrix completion tasks where it can learn from historical data and predict unknown entries in the matrices. The latent factors are first learnt through only the known entries in the matrices, that can be considered as the training data. Completed matrices, obtained by multiplying the learnt latent factors, includes the predictions for the unknown entries. This is similar to the setting used in recommendation tasks, which has also been used in other bioinformatics applications, e.g., in (Natarajan and Dhillon, 2014).

### Limitation of CMF

If multiple input matrices to CMF contain the same row and column entity-types, then CMF (or gCMF) cannot learn a unique representation for each entity. For instance, consider two matrices with pairwise gene coexpression (*X*_1_) and mutual exclusivity information (*X*_2_). where all the row and column entities are genes. But it is impossible to reconstruct two different matrices, such that, 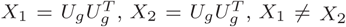, using unique latent factors *U*_*g*_ for genes (*g*). The same problem occurs if the row and column entities are identical in two or more input matrices. E.g., matrices containing gene expression (*X*_1_) and copy number alteration (*X*_2_), have relations between entities gene (*g*) and patients (*p*) and it is impossible to reconstruct the matrices 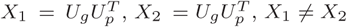, using unique latent factors *U*_*g*_, *U*_*p*_.

## 4 Our Approach

To model collections of matrices that may have multiple matrices with identical row and column entities, we propose three solutions. The first two solutions rely on a transformation before CMF can be applied. The third solution modifies the model to use a matrix-specific factor to directly learn latent representations from the input data. They can all be viewed as different forms of link functions in the formulation of Singh and Gordon (2008); the difference is that in our case, the link function is matrix-specific.

### Transformation Using PCA

We use Principal Components Analysis (PCA) to obtain eigenvectors of the matrix. The leading eigenvectors can be selected as the columns in the transformed matrix. This also allows us to reduce dimensionality of the matrix, if required. We choose the minimum number of principal components required for the cumulative explained variance ratio to be greater than 0.9. Thus, we can transform a matrix with identical row and column entities (say, *e*_1_) to a matrix where the row entity is *e*_1_ and column entity is features that are matrix-specific, as shown in fig. 3a. Such features have been found to be effective in other matrix completion based predictive models, e.g., in (Natarajan and Dhillon, 2014). When this transformation is applied before using CMF or gCMF, we call the method **pca-CMF** or **pca-gCMF**, respectively.

**Fig. 3:**
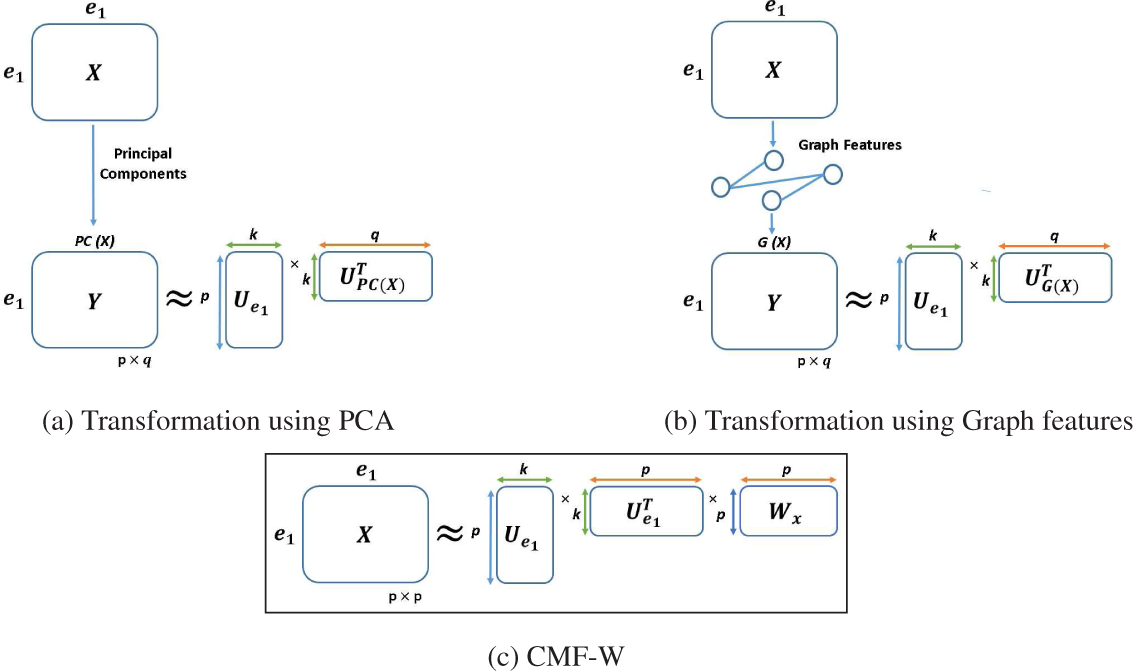
Overview of our solution. Multiple matrices with identical row and column entities in an input collection of matrices cannot be handled by CMF directly. If PCA (fig. 3a) or graph-based features (fig. 3b) are used to transform such matrices, then CMF can be applied on the transformed matrices, since the column entities in each of the transformed matrices are different and not identical to their row entities. Our model CMF-W (fig. 3c) extends CMF by using a matrix-specific weight (W) that can model different data in each input matrix with identical row and column entities.

### Transformation Using Graph Features

Matrices with identical row and column entity-types can be viewed as adjacency matrices of graphs where each entity is used to form the node set. When the row and column entity-types are not identical, the matrix can be viewed as an adjacency matrix of a bipartite graph, with the row entities being one set of nodes and column entities being the other set of nodes. In both cases, a cell entry can be considered to be an edge label. We can transform the adjacency matrix to another matrix where the column entity is formed by graph-based features that are matrix-specific, as shown in fig. 3b. To transform the input matrices, we use graph features that were found to be useful in predicting SL interactions in yeast using PPI networks (Paladugu *et al.*, 2008). These include the node degree, closeness centrality, betweenness centrality (Freeman, 1977), information centrality (Stephenson and Zelen, 1989), eigenvector centrality (Bonacich, 1972), Gil-Schmidt Power Index (Gil-Mendieta and Schmidt, 1996), and the Flow Betweenness Score (Freeman *et al.*, 1991). Appendix A has definitions of these features. When this transformation is applied before using CMF or gCMF, we call the method **gr-CMF** or **gr-gCMF**, respectively.

### CMF-W

In this approach, we modify the CMF model by incorporating matrix-specific weights to handle matrices with identical row and column entity types. Each such matrix is modeled as a product of three factors:

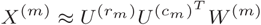

The first two factors are the same as in CMF, i.e., the row and column entity representations, while the third factor is a matrix-specific weight *W* ^(*m*)^. This third factor models the (unknown) transformation in each input source responsible for different values and datatypes. Thus, for a matrix *X*^(*m*)^ with identical row and column entities ((*r*_*m*_) = (*c*_*m*_) = (*g*_*m*_)), we have 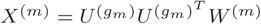, where *W* can be different for two matrices with identical row and column latent factors (fig. 3c). The latent factors are learned by solving the optimization problem:

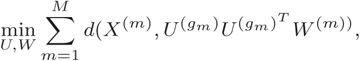

where *d* is the Frobenius norm of the difference between *X*^(*m*)^ and 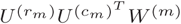. For *m* × *n* matrix *X*^(*m*)^ and latent dimension *k*, the dimensions of 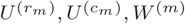 are *m* × *k, n* × *k* and *n* × *n* respectively. We use the Adam optimization algorithm (Kingma and Ba, 2015) to solve the optimization problem.

## 5 Experiments

We pose the problem of SL prediction as a binary classification task on pairs of genes, with positive class indicating SL interactions and negative class indicating no SL interactions.

### Baselines

We use five state-of-the-art methods designed for predicting SL pairs: ISLE, DAISY, MetaSL, MNMC and MCA. These methods have been tested with different input datasets. Our first three experiments match the input data used in MCA, DAISY and ISLE respectively. We conduct a fourth experiment with another dataset. Details are given below.

### Evaluation Metric

We use two metrics to evaluate performance. The first is AUC (Area under the ROC Curve), on held-out test sets, that has been used in all the baselines that we compare with. However, while there is previous evidence of positive SL pairs (e.g., through knock-out screens), the evidence for negative pairs is weaker and so, these pairs could be considered as unlabelled. Further, in the application of SL prediction, it is more important to penalize false positives than false negatives. So, our second metric is ‘probability-at-n’, that is used in positive-unlabelled learning and in similar applications, e.g., gene-disease prioritization (Natarajan and Dhillon, 2014). For the *i*^*th*^ gene, we order the other genes (indexed by *j*) by scores assigned by the predictive models. For every gene pair (*i, j*) in the held-out test set we record the rank of the gene *i* in the list associated with gene *j*. Probability-at-n is the probability that the rank (at which an SL pair is retrieved) is less than a threshold *n* (i.e., the cumulative distribution of the ranks). This measures the probability of recovering a true SL interaction in the top-n predictions for a given gene. A small value of *n* is desirable and we report results for *n* ≤ 180.

### Experiment 1

We first compare our method with that of Lu *et al.* (2015), on their published datasets. They obtain 270 SL pairs and 5660 non-SL pairs from two previous studies (Laufer *et al.*, 2013; Vizeacoumar *et al.*, 2013). Using Copy Number Variation (CNV) data and RNAseq data they design five features for each gene pair based on homozygous, heterozygous or mixed co-loss events, likelihood of simultaneous under-expression and likelihood of inverse expression (i.e., when one gene is over-expressed and the other under-expressed). We can represent these features as five matrices with genes as both row-entity and column-entity in each matrix and where, the *ij*^*th*^ cell contains the feature value for the gene pair (*i, j*).

Due to the imbalance present in the data (only 4.6% data in the positive class of SL pairs), we follow an undersampling based approach similar to that of Lu *et al.* (2015). We conduct 10 experiments; retaining the same 270 SL pairs in each experiment, 270 non-SL pairs are randomly sampled from the 5660 non-SL pairs independently for each experiment. Then, in each experiment, we randomly select 70% of the SL pairs (378 pairs) for training and remaining 30% (162 pairs) as the test set. The average performance over these 10 experiments is reported. On this dataset, we compare the performance of MNMC, MCA and MetaSL with our CMF-based approaches.

### Experiments on Breast Cancer Data

In the remaining three experiments, we use all 245 SL pairs associated with breast cancer as reported in SynLethDB (Guo *et al.*, 2015). Let *S* be the set of genes in these pairs. Pairs in the negative samples, i.e., pairs that are not SL, may have a gene that can be an SL partner (with some other gene) or may have both genes that are not involved in any known SL interactions. To test both these cases, we select negative samples, denoted by *N*, from the HGNC database (Bruford *et al.*, 2007) after excluding genes reported in any SL interaction in SynLethDB and those reported to be essential in (Vizeacoumar *et al.*, 2013; Marcotte *et al.*, 2012). We construct our negative samples by randomly selecting 200 pairs (*g*_*i*_, *g*_*j*_) such that *g*_*i*_ ∈ *S, g*_*j*_ ∈ *N* and 45 pairs such that *g*_*i*_ ∈ *N, g*_*j*_ ∈ *N*. Thus, there are a total of 332 unique genes used and 490 labelled pairs. We call this matrix the *SL-label* matrix in the following. See Appendix B.1 for more details and a schematic of our matrix.

We use 3-fold cross validation to evaluate and compare performance of various methods. In addition, we also perform stratified 3-fold cross validation, where the proportion of positive and negative class samples are balanced across the folds. In the case of methods that are not based on machine learning, such as DAISY or ISLE, the training data in each fold is not utilized and predictions are made directly on the test data in each fold. The statistical tests in DAISY and ISLE are specific to their input data, and so, these results are only shown for experiments 2 and 3 respectively. For MCA, MNMC and MetaSL, all the input matrices, except SL-label, are concatenated and used as features in each experiment. The average probability-at-n and average AUC (with standard deviation) across the folds, for all the methods are reported.

### Experiment 2

We compare our methods with DAISY (Jerby-Arnon *et al.*, 2014), using matched data sources. DAISY conducts three independent statistical tests using Somatic Copy Number Alteration (SCNA), mutation profiles (containing information of deleterious mutations, i.e., whether a gene has frameshift or nonsense mutations), gene essentiality profiles, and pairwise gene co-expression data. We obtained SCNA, mRNA gene expression data and mutation profiles for breast cancer patients in TCGA (TCGA, 2012) using cBioPortal (Gao *et al.*, 2013; Cerami *et al.*, 2012) and Firehose. Essentiality profiles are based on those curated in (Marcotte *et al.*, 2012) for breast cancer in addition to the (∼ 16,000 essentiality) genes listed in (Vizeacoumar *et al.*, 2013).

In DAISY a pair is predicted to be SL if it passes all three tests. We denote the first test by DAISY-1, and the method comprising the first and third test is called DAISY-3. The second test is not included because in our experiments, no gene pairs were selected after the second test. Similar results were observed by Jerby-Arnon *et al.* (2014) who report that the shRNA-based functional examination, i.e., the second test, is not predictive on its own (with an AUC of 0.477 in their larger dataset). They also use the second test only as a soft constraint after identification of gene pairs using the first and third test. For CMF-based methods, we use four matrices in addition to the SL-label matrix: SCNA, gene expression data, essentiality profile and pairwise co-expression data. Since both SCNA and gene expression data have the same row-entity (gene) and column entity (patient), we chose one of the matrices, SCNA, for (graph and PCA) transformations and retained the other, gene expression, without any transformation. Implementation details of DAISY and our approach are described in Appendix B.2.

### Experiment 3

To compare with ISLE, we use the software and data provided by them (Lee *et al.* (2018)), using only the breast cancer data. We obtain phylogenetic similarity using the phylogenetic profile database (Sadreyev *et al.*, 2015). For our CMF-based methods we use the scores for 86 species provided by the database directly.

### Experiment 4

We also compare the performance of our methods on another dataset where we use features for each pair of genes, derived from five sources: Co-expression from StringDB (Szklarczyk *et al.*, 2014); Mutual Exclusivity scores for breast cancer from TiMEx (Constantinescu *et al.*, 2015); Pathway Co-membership, using the Canonical pathway database from Broad Institute’s Molecular Signatures Database (MSigDB) (Subramanian *et al.*, 2005); Protein Complex Co-membership, using the CORUM protein complex database (Giurgiu *et al.*, 2018); and Protein-Protein Interactions (PPI) scores from the Hippie database (Alanis-Lobato *et al.*, 2017). In the two co-membership matrices, we assign a 1 to a gene pair if they belong to the same pathway or protein complex, otherwise a 0. All the six matrices have genes as row and column entities and are of dimensions 332 × 332.

All the matrices used in experiments 2,3 and 4 in our CMF-based approaches are shown in table 1. Note that although the input features differ across these three experiments, the test set (i.e., held-out entries in the SL-label matrix) across the three folds are identical in these three experiments and hence, the results are comparable.

**Table 1.** Input matrices, their row and column entities and dimensions in our methods in experiment 2 (left), experiment 3 (middle), experiment 4 (right).

| Matrix | Row Entity | Col Entity | Row Dim | Col Dim | Matrix | Row Entity | Col Entity | Row Dim | Col Dim | Matrix | Row Entity | Col Entity | Row Dim | Col Dim |
| --- | --- | --- | --- | --- | --- | --- | --- | --- | --- | --- | --- | --- | --- | --- |
| SL-label | Gene | Gene | 332 | 332 | SL-label | Gene | Gene | 332 | 332 | SL-label | Gene | Gene | 332 | 332 |
| Essentiality Profile | Gene | Gene | 332 | 332 | Essentiality Profile | Gene | Gene | 332 | 332 | Co-expression | Gene | Gene | 332 | 332 |
| mRNA Gene Expression | Gene | Patient | 332 | 1075 | mRNA Expression | Gene | Patient | 332 | 1075 | Mutual Exclusivity | Gene | Gene | 332 | 332 |
| SCNA Level | Gene | Patient | 332 | 1075 | SCNA Level | Gene | Patient | 332 | 1075 | Pathway Co-membership | Gene | Gene | 332 | 332 |
| Pairwise co-expression | Gene | Gene | 332 | 332 | Phylogenetic Score | Gene | Species | 332 | 86 | Protein Complex Co-membership | Gene | Gene | 332 | 332 |
|  |  |  |  |  |  |  |  |  |  | Protein-Protein Interaction (PPI) | Gene | Gene | 332 | 332 |

## 6 Results

Fig. 4 (leftmost) shows the performance of all our methods and three ensemble-based methods on the published dataset of Lu *et al.* (2015) comprising five evolutionary features for each pair of genes. CMF-W, pca-CMF and gr-CMF do not outperform previous methods MCA, MetaSL and MNMC. However, pca-gCMF and gr-gCMF significantly outperform all other methods, both achieving average AUC of more than 0.9. The same trend is observed with respect to probability-at-N. At all values of N, pca-gCMF and gr-gCMF outperform MCA, MetaSL and MNMC. The performance of CMF-W is comparable to the baselines.

**Fig. 4:**
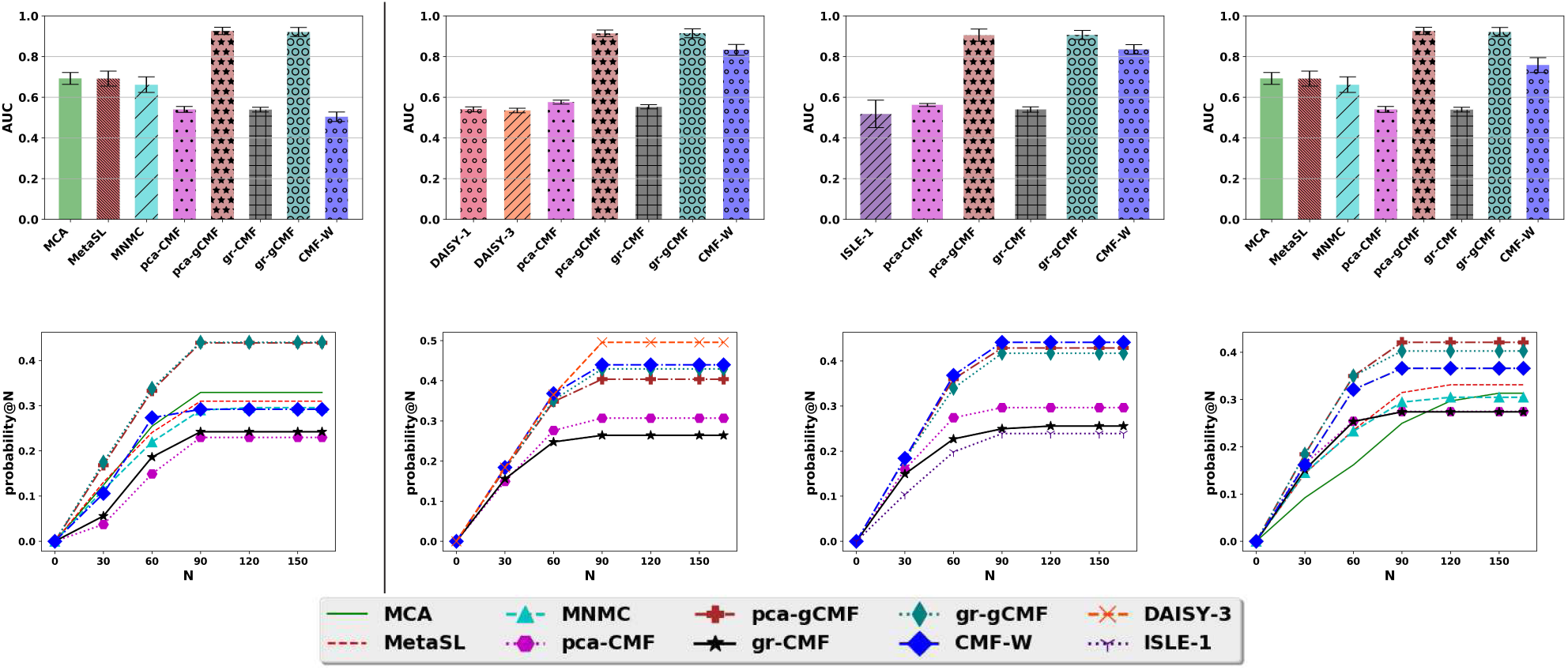
Results on 3-fold CV. Columns (left to right): Experiments 1–4 (identical test sets in 2–4). Rows: AUC (above), Probability-at-N (below).

The second column of fig. 4 compares the performance of our CMF-based methods with DAISY in experiment 2. The AUC achieved by DAISY is lower than the reported AUC in (Jerby-Arnon *et al.*, 2014). Although we have used the same data sources, our test sets are different and restricted to breast cancer only. Similar low AUC for DAISY are reported in other datasets (e.g. (Lee *et al.*, 2018)). While pca-CMF and gr-CMF have AUC comparable to that of DAISY, pca-gCMF, gr-gCMF and CMF-W outperform DAISY. However, with respect to probability-at-N, DAISY, CMF-W and pca-gCMF are comparable (and better than the rest) for *N* ≤ 60, and DAISY outperforms all the methods at *N* ≥ 60. With respect to CMF-based approaches, DAISY has comparable (at *N* ≤ 60) or better (at *N* ≥ 60) sensitivity while its specificity may be lower resulting in lower AUC.

The third column of fig. 4 compares the performance of our CMF-based methods with ISLE in experiment 3. None of the pairs passed the second test and so we show the results only for ISLE-1. CMF-W, pca-gCMF and gr-gCMF outperform the remaining methods that have comparable AUC. A similar trend is seen in prbability-at-N values with ISLE underperforming at all values of *N*. In the last column of fig. 4, the AUC of MCA, MetaSL and MCMC is found to be better than those of pca-CMF and gr-CMF. However, pca-gCMF, gr-gCMF and CMF-W outperform other methods in experiment 4 in both AUC and probability-at-N.

Note that in experiments 2, 3 and 4, the test sets used are identical across the folds. Hence these results are comparable. In general, we observe that that performance of gCMF and CMF-W is consistently better than that of CMF. Experiments 2,3 and 4 with stratified 3-fold CV are discussed in Appendix C, where we observe the same same performance trends. Experiment 4 is also conducted with four other random samples of the negative set *N*; these results, discussed in Appendix D also show the same performance trends. For all the CMF-based approaches, we repeat the experiments with different values (2, 5, 10, 50) of the latent dimension *K*. The best performance is seen for *K* = 2 with results deteriorating slightly with increasing *K* (shown in Appendix F). We investigate this further for pca-gCMF and observe that the distibution of latent factor values are more peaked at *K* = 2 and more flat at *K* = 50 (results in Appendix G). Thus, more sparse solutions are correlated with better performance in gCMF. This is also observed in the difference of performance between pca-CMF and pca-gCMF (or gr-CMF and gr-gCMF) with the latter, that yields sparse solution, outperforming the former in all our experiments. Better performance of CMF-W over CMF, can be attributed to better optimization method (Adam) used in CMF-W as well as better modeling of matrix-specific parameters (*W*).

An advantage of our CMF-based approach is that it can be used with arbitrary collections of matrices. This can be used to investigate the relative value of the ‘signal’ provided by each data source or combinations of data sources by systematically using subsets of data matrices for prediction. Such an analysis is described in Appendix E that shows the relative importance of each data matrix for experiments 2–4.

## 7 Conclusion

Integration of data from heterogeneous sources is a key challenge in bioinformatics, particularly in cancer studies. Collective Matrix Factorization (CMF) and its variants can model heterogeneous data, represented as relations between entities in matrices. However, CMF cannot be used directly when two or more matrices in the input have the same row and column entities, a case that is commonly found in biological datasets (and in all the datasets in our experiments). By overcoming this limitation, our techniques can be effectively utilized on many large heterogeneous datsets. We illustrated the advantage of our methods in predicting synthetic lethality in gene pairs using a machine learning based matrix completion approach on four different datasets.

Previous methods for predicting SL pairs, like DAISY and ISLE, use statistical inference tests that are specifically designed for the input data they use. More general machine learning approaches, like MCA, require considerable effort in feature engineering to obtain features with high predictive value. In contrast, our approach can directly use relations of genes with other entities like patients or species, and could also benefit from auxiliary data sources containing different but related entities (e.g., patients and their clinical features). Thus, our approach can seamlessly integrate multiple heterogeneous data sources, which can be either specific features (in experiments 1, 2 and 3) or those derived, with considerably less feature engineering, from multiple existing databases (experiment 4). In fact, the versatility and accuracy of our method is best indicated by comparing its performance across experiments 2,3 and 4 (that use the same test data). Our approach achieves the highest AUC in experiment 4, without the complex feature engineering used in experiments 2 and 3.

Future work can further extend the modeling capability of these methods, and evaluate the methods on other datasets, including other applications that can benefit from integrating heterogeneous data sources. Strategies to improve learning, e.g., through better initialization, can be explored. We also plan to validate our predictions for previously untested gene pairs through CRISPR screens.

## Appendix

### A Graph Features used in gr-CMF and gr-gCMF

- **Degree**. In undirected networks, the node degree of a node *v* is the number of edges linked to *v*. A self-loop of a node yields a degree of 2. The node degrees measures the number of direct interactions in the network.
- **Closeness centrality**. It is a measure of centrality of a node. For a node *x*, it is given by 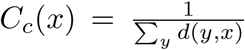, where *d*(*y, x*) is the distance of the shortest path between vertices *x* and *y* (Sabidussi, 1966). The normalized version has a multiplicative factor equal to the number of nodes in the network.
- **Betweenness centrality**. Another centrality measure, the betweenness centrality of a node *v* is given by: 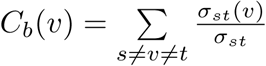, where *σ*_*st*_ is the total number of shortest paths from node *s* to node *t* and *σ*_*st*_ (v) is the number of those paths that pass through *v* (Freeman, 1977).
- **Information centrality**. Let *A* be an adjacency matrix of a network, *D* a diagonal matrix of the degree of each node and *J* a matrix with all its elements equal to one; we define *B* = *D* – *A* + *J* and let *C* = *B*^−1^. This yields the information matrix I with elements *I*_*vj*_ = (*C*_*vi*_ + *C*_*jj*_ + *C*_*vj*_). The information centrality *IC*(*v*) of node *v* is then defined as harmonic mean: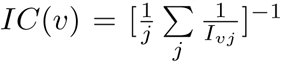, where the information measure *I*_*vj*_ between nodes is defined as the reciprocal of the topological distance *d*_*vj*_ between the corresponding nodes, 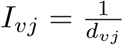(Stephenson and Zelen, 1989).
- **Eigenvector centrality** The eigenvector centrality of a node *v* is defined as the *v*^*th*^ element of the principal eigenvector of the adjacency matrix. This principal eigenvector is normalized such that its largest entry is 1 (Bonacich, 1972).
- **Gil-Schmidt Power Index**. This index generalizes degree centrality by taking into account not just the order of the neighborhood set of the node, but also a weighted sum of the orders of each *kth*-neighborhood set in the network with respect to the indexed node Gil-Mendieta and Schmidt (1996).
- **Flow Betweenness Score**. The flow betweenness of a vertex, *v* is defined by: 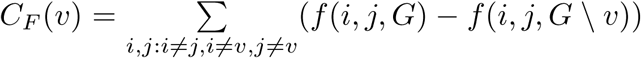, where *f* (*i, j, G*) is the maximum flow from *i* to *j* within *G*. Intuitively, it is the total flow mediated by *v* (Freeman et al., 1991).

Typically, data matrices, and hence the corresponding graphs, in bioinformatics are not sparse and so, we can use a threshold value on the entries to induce sparsity. E.g., we can construct a graph, without edge labels, by considering all cells with values greater than the threshold. We choose a threshold value of 0 for our experiments.

### B. Experiment Settings

#### B.1 SL Label Matrix

For experiments 2,3 and 4, we use all 245 SL pairs associated with breast cancer as reported in SynLethDB (Guo et al., 2015). Let *S* be the set of genes in these pairs. Pairs in the negative samples, i.e., pairs that are not SL, may have a gene that can be an SL partner (with some other gene) or may have both genes that are not involved in any known SL interactions. To test both these cases, we select negative samples in the following manner. From an initial set of 41,289 genes in the HGNC database (Bruford et al., 2007), we exclude those genes that are reported in any SL interaction in SynLethDB (5,131 genes) and also exclude those genes reported to be essential in both (Vizeacoumar et al., 2013) and (Marcotte et al., 2012) (15,715 genes). In total, there are 25,388 (unique) genes after the exclusions, that we denote by set *N*. We construct our negative samples by randomly selecting 200 pairs (*g*_*i*_, *g*_*j*_) such that *g*_*i*_ ∈ *S, g*_*j*_ ∈ *N* and 45 pairs such that *g*_*i*_ ∈ *N, g*_*j*_ ∈ *N*. Thus, there are a total of 332 unique genes used and 590 labelled pairs. See figure B.1 for a schematic of our matrix, the diagonal indicating that the matrix is symmetric. We call this matrix the *SL-label* matrix.

**Figure B.1:**
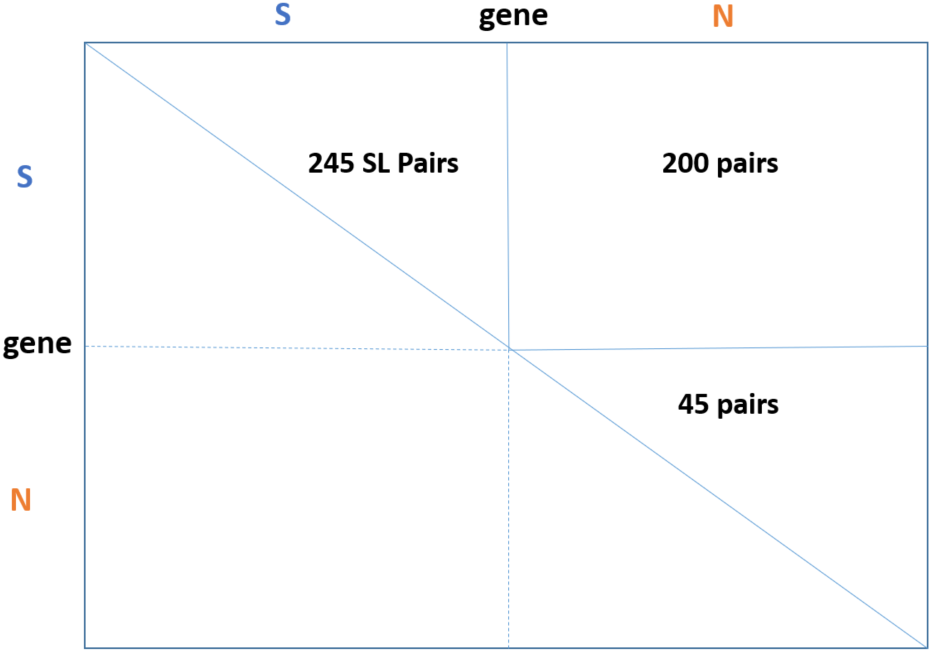
Schematic of SL-label Matrix with curated SL and non-SL interactions used in experiments 2,3,4. Entries are symmetric across the diagonal.

#### B.2 Settings for Experiment 2

To compare our methods with DAISY (Jerby-Arnon et al., 2014), we use the same data sources. DAISY conducts three independent statistical tests using Somatic Copy Number Alteration (SCNA), mutation profiles (containing information of deleterious mutations, i.e., whether a gene has frameshift or nonsense mutations), gene essentiality profiles, and pairwise gene coexpression data. We obtained SCNA, mRNA gene expression data and mutation profiles for breast cancer patients in TCGA (TCGA, 2012) using cBioPortal (Gao et al., 2013; Cerami et al., 2012) and Firehose ^1^. Essentiality profiles are based on those curated in (Marcotte et al., 2012) for breast cancer in addition to the (∼ 16,000 essentiality) genes listed in (Vizeacoumar et al., 2013).

In DAISY a pair is predicted to be SL if it passes all three tests. For our analysis, we check the results in a cumulative manner, as described below, to obtain three results. Following Jerby-Arnon et al. (2014), we consider a gene to be inactive in a sample if it is underexpressed and its SCNA is below −0.3 or if it is mutated with a deleterious mutation; a gene is considered to be overactive in a sample if it is overexpressed and its SCNA is above 0.3. A gene is defined as under-expressed in a given sample if its expression level is below the 10th percentile of its expression levels across all samples in the data set or its SCNA is below −0.3 or if it is mutated with a deleterious mutation. Similarly, a gene is over-expressed if its expression level is above its 90th percentile or its SCNA is above 0.3.

For the first test, that we call DAISY-1, a Wilcoxon rank sum test is used to check if, for a pair (A,B), gene B has a significantly higher SCNA level in samples in which gene A is inactive (overactive) than in the rest of the samples (and similarly, for the pair (B,A)). Gene pairs that pass the test (p-value < 0.05 following Bonferroni correction for multiple hypotheses testing) are predicted to be SL. For the second test, since we did not have access to shRNA-based functional screen data, we checked for essentiality using the data from (Marcotte et al., 2012). For a pair of genes (A, B), we conduct a Wilcoxon rank sum test to check if gene B is significantly more essential in samples in which gene A is inactive (overactive) than in the rest of the samples (similarly, for (B,A)). We denote by DAISY-2, the method that conducts both the first and the second test. For the third test, we consider a pair of gene to be positively correlated if it is significantly positively correlated in at least one of 7 transcriptomic datasets (containing gene expression profiles for the following cancers from TCGA : Breast Cancer, Colon Cancer, Colorectal Cancer, Glioblastoma, Liver, Lung and Ovarian Cancer). Correlation is measured by Spearman’s correlation coefficient following Bonferroni correction for multiple hypotheses testing. We denote by DAISY-3, the method that conducts all three tests. None of the pairs passed the second test and so we show the results only for DAISY-1 and DAISY-3.

For CMF-based methods, we use four matrices in addition to the SL-label matrix: SCNA, gene expression data, essentiality profile and pairwise co-expression data. Since both SCNA and gene expression data have the same row-entity (gene) and column entity (patient), we chose one of the matrices, SCNA, for transformation in CMF and retained the other, gene expression, without any transformation.

We obtain a binary matrix from the pairwise co-expression data using the test for positive correlation in DAISY-3. We obtain a binary matrix from the gene-expression profile, with a 1 if a given sample is under-expressed (below its 10th percentile of its expression level across or all samples) or over-expressed (above its 90th percentile), otherwise a 0 for each pair of genes. We also obtain a binary matrix from the SCNA profile, with a 1 if a given sample is below −0.3 or above 0.3, otherwise a 0 for each pair of genes.

### C Stratified 3-fold CV results

**Figure C.1:**
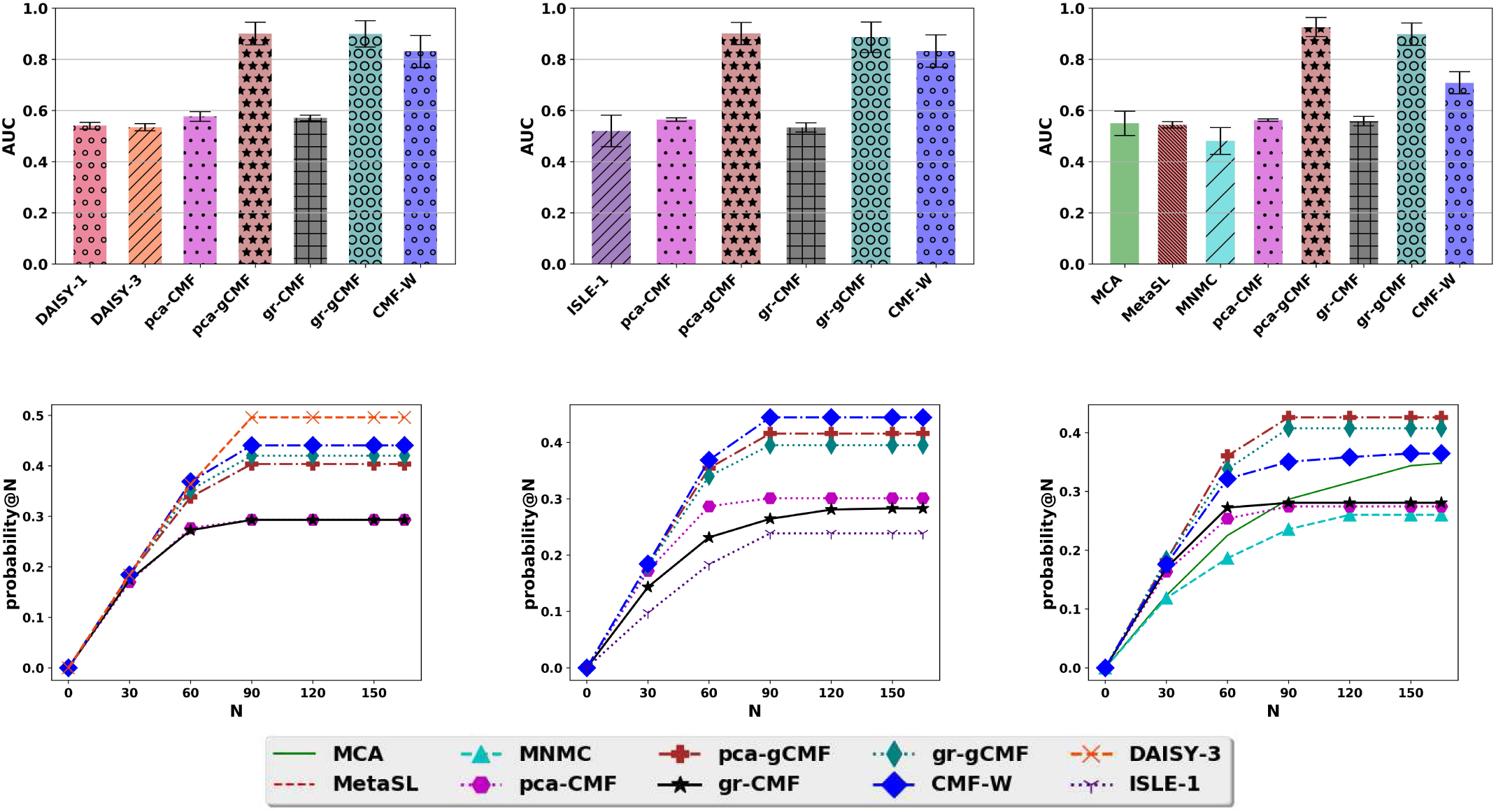
AUC (row above) and Probability-at-N (row below) averaged over 3-fold CV for experiments 2–4 (columns left to right).

Figure C.1 shows the results of experiments 2-4 as discussed in section 5 with stratified 3-fold CV.

### D Negative Samples

**Figure D.1:**
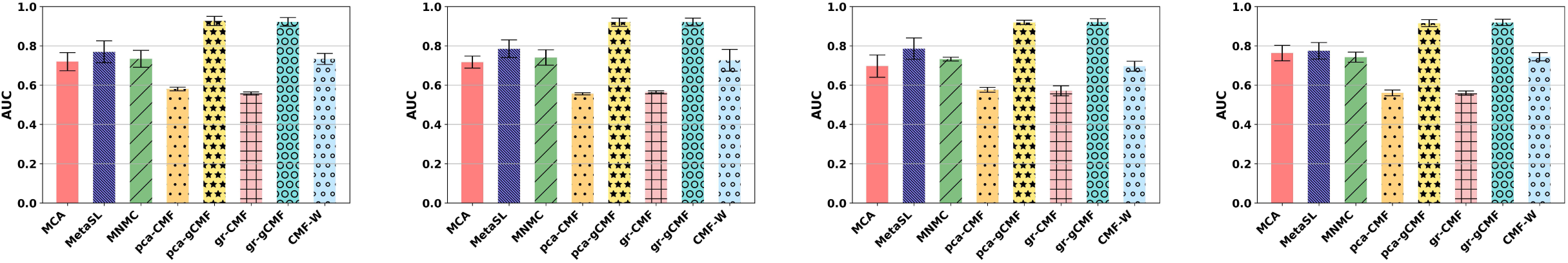
AUC averaged over 3-fold CV (with standard deviation) for four other randomly sampled negative sets *N* in experiment 4.

Figure D.1 shows AUC averaged over 3-fold CV (with standard deviation) for four other randomly sampled negative sets *N* in experiment 4. There is no significant change in performance trends across the negative samples. PCA-gCMF and gr-gCMF has the best performance across all five random selections.

### E Feature Selection

**Figure E.1:**
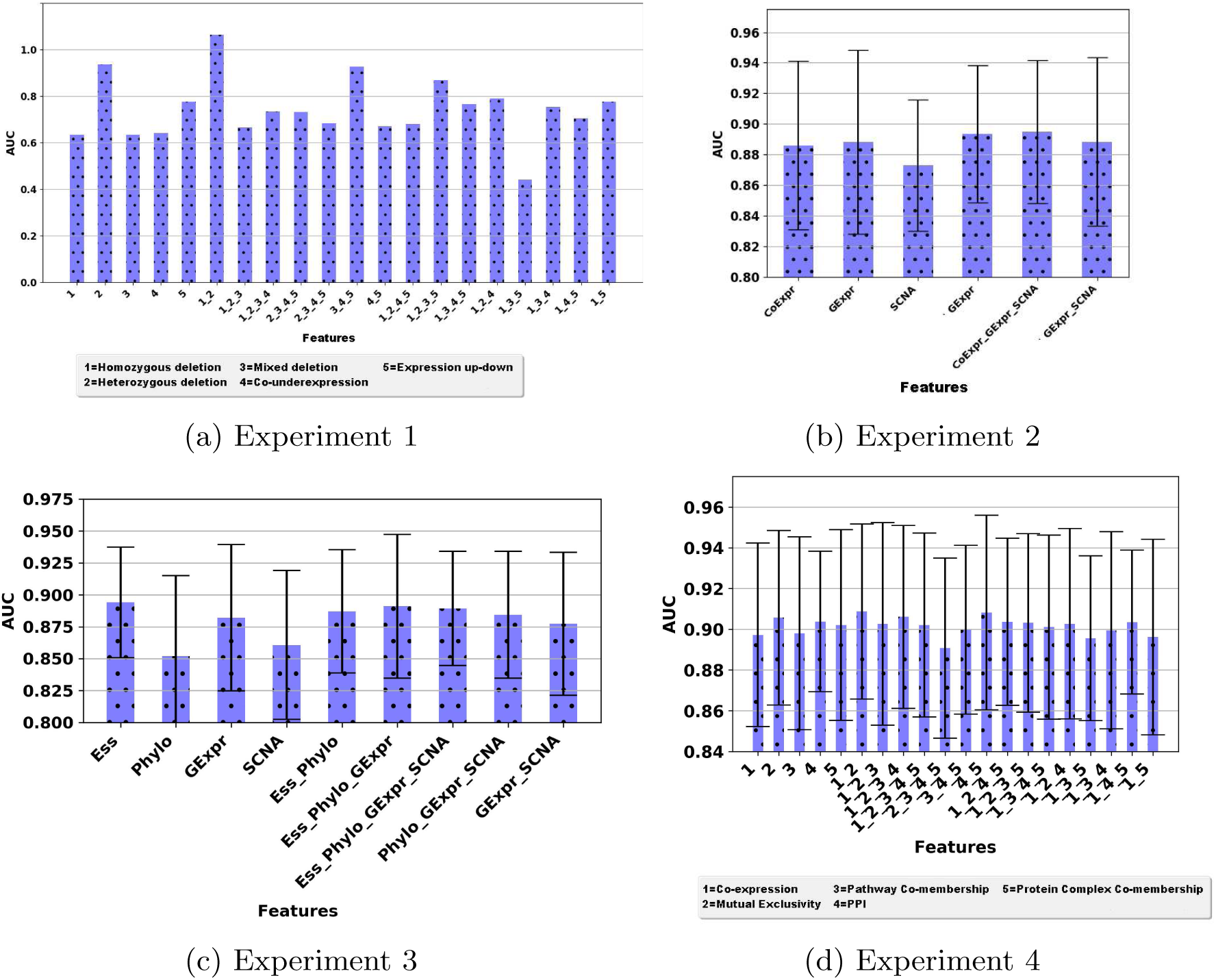
Predictive signal from each data source and combinations of data sources in experiments 1–4.

An advantage of our CMF-based approach is that it can be used with arbitrary collections of matrices. This can be used to investigate the relative value of the ‘signal’ provided by each data source or combinations of data sources by systematically using subsets of data matrices for prediction. Figure E.1 shows the results of such an analysis for experiments 1–4. In each experiment we used each data source individually to predict SL pairs and then used each combination of data sources (all subsets) to predict SL label using pca-gCMF. Note that the SL-label matrix that has the known SL labels and unknown entries that are predicted is used in all the cases.

In experiment 1 (figure E.1a), we observe that the highest AUC is achieved by the combination of feature matrices 1 and 2 (more than using all the five matrices). In experiment 2 (figure E.1b), using SCNA matrix alone has higher AUC than using all the three matrices. In experiment 3 (figure E.1c), use of all four matrices has the highest AUC with low standard deviation. In experiment 4 (figure E.1d), any of the matrices or combinations of them have roughly equivalent predictive signal.

### F Selecting K

**Figure F.1:**
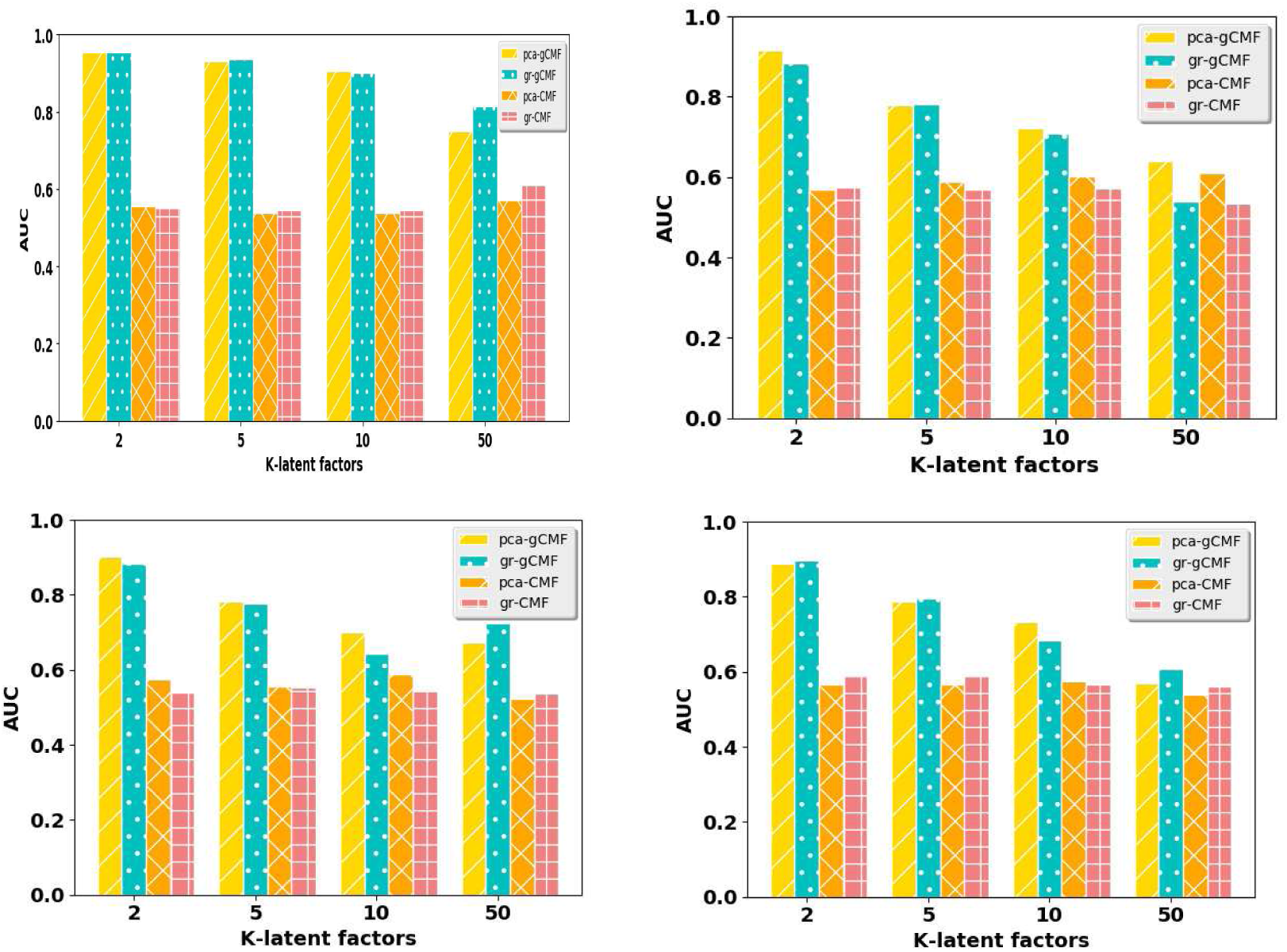
AUC values at different values of latent dimension *K* for experiment 1 (top left), experiment 2 (top right), experiment 3 (bottom left) and experiment 4 (bottom right) for our CMF-based approaches.

We empirically investigate the effect of different choices of latent dimension *K* in our CMF-based approaches. Figure F.1 shows the average AUC obtained by our transformation-based approaches for four different choices (2, 5, 10, 50) of *K* in experiments 1–4.

CMF is less sensitive to the choice of *K* and we observe roughly the same performance at all four values of *K*. However the AUC obtained by CMF is lower than that of gCMF at all the values. gCMF is more sensitive to the choice of *K* with the performance decreasing with increasing value of *K*. The best AUC values are obtained at *K* = 2. The same trends are observed with both – PCA and graph-based – transformations.

### G Sparsity levels

Tables G.1, G.2, G.3 and G.4 show the distribution of the values (in 10 bins after min-max scaling) in the learnt latent factors in experiments 1–4 respectively for four choices of *K* : 2, 5, 10, 50. The distributions of the values in the latent factors for ‘gene’ entity (for all four values of *K*) are shown in figures G.1, G.2, G.3 and G.4 respectively for experiments 1–4. All these results are for PCA-gCMF only.

We observe that the distributions are more peaked at *K* = 2 and more flat at *K* = 50. This indicates that at lower values of *K* more values are concentrated in fewer bins compared to those in higher values of *K*. The performance trends seen in figure F.1, and these distributions suggest that more sparse solutions are correlated with better performance in gCMF. This is also observed in the difference of performance between pca-CMF and pca-gCMF (or gr-CMF and gr-gCMF) with the latter, that yields sparse solution, outperforming the former in all our experiments.

**Figure G.1:**
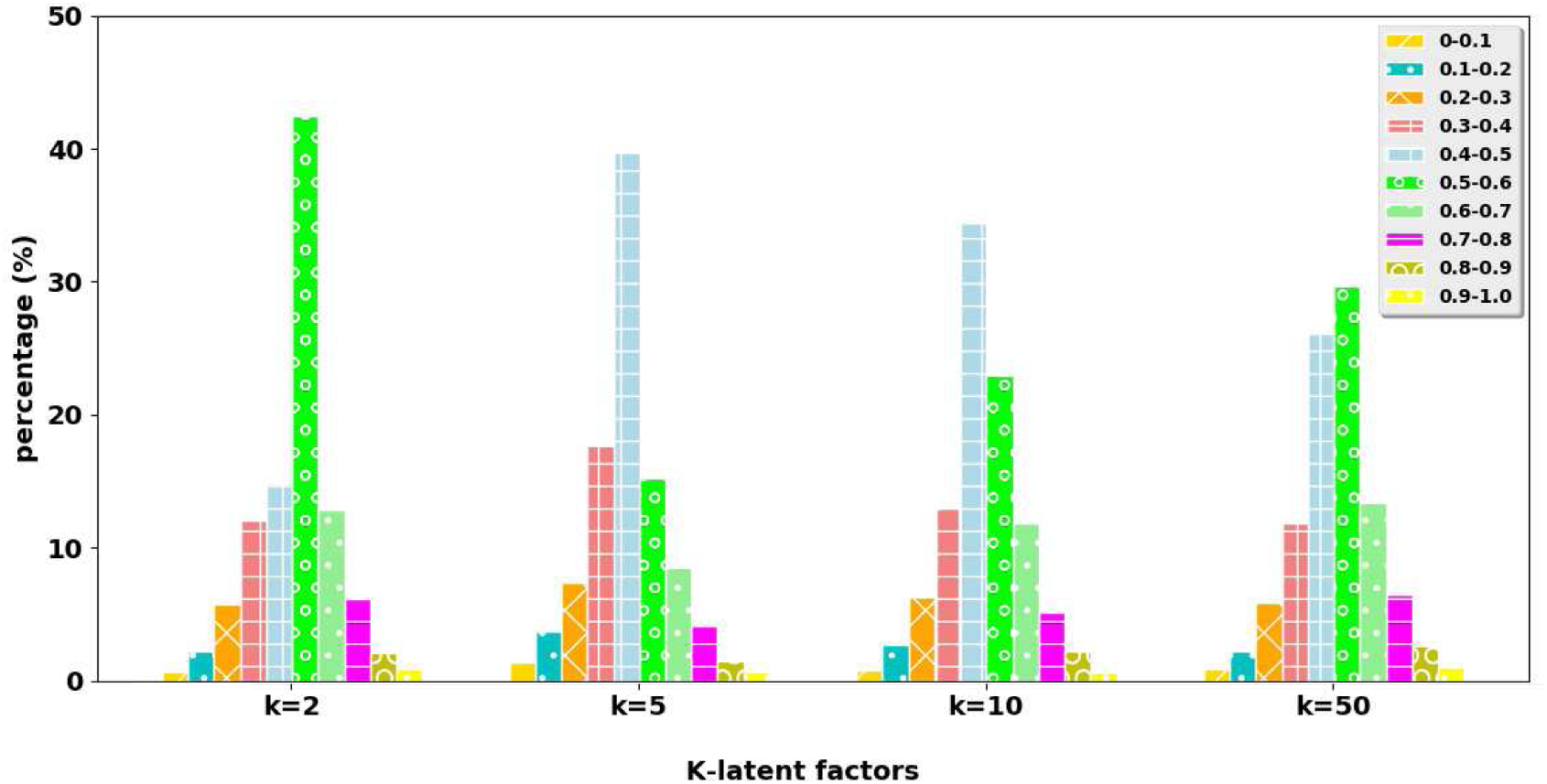
Experiment 1: distribution of values (in 10 bins after min-max scaling) of gene latent factor at four choices of *K*.

**Figure G.2:**
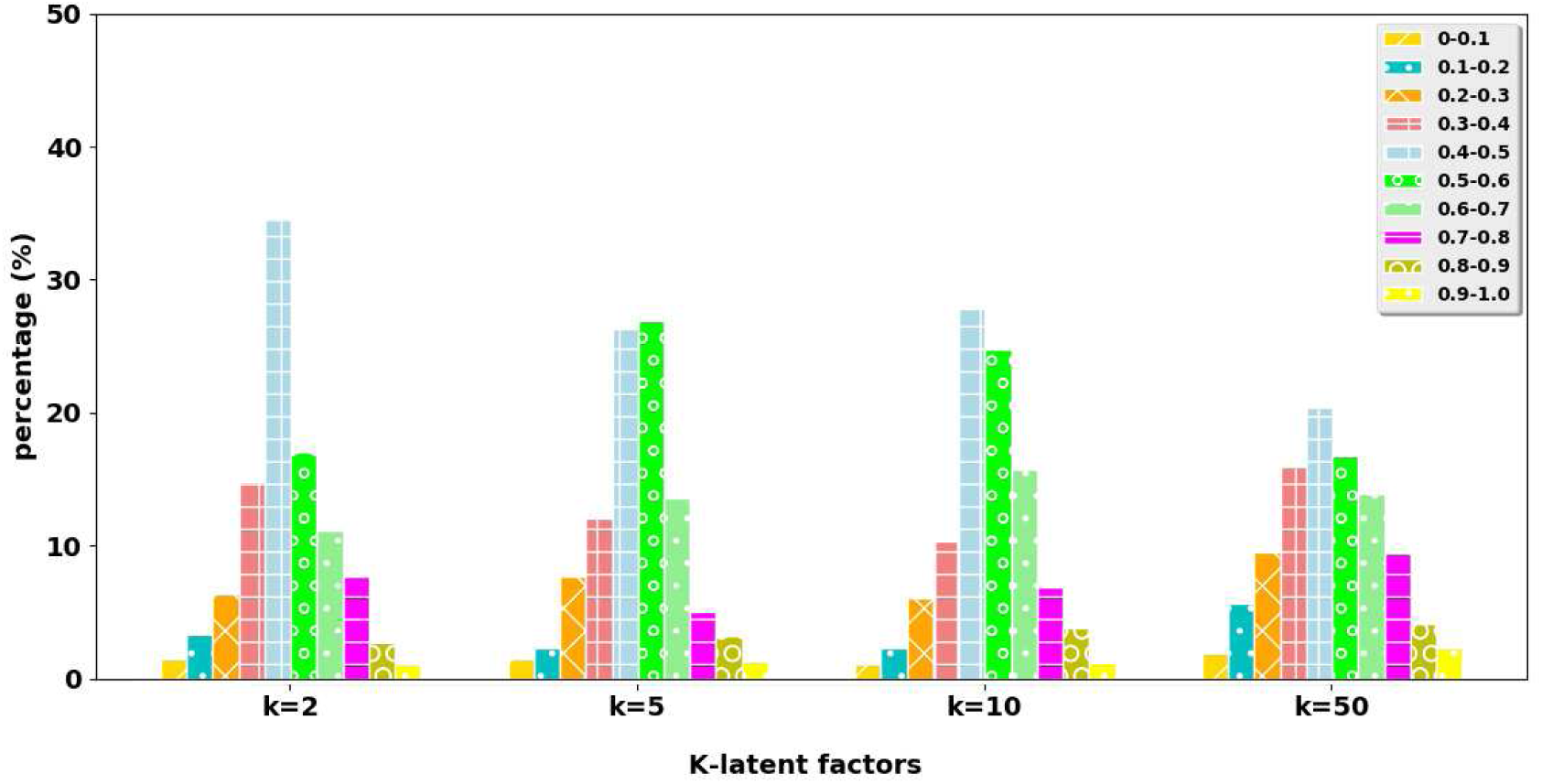
Experiment 2: distribution of values (in 10 bins after min-max scaling) of gene latent factor at four choices of *K*.

**Figure G.3:**
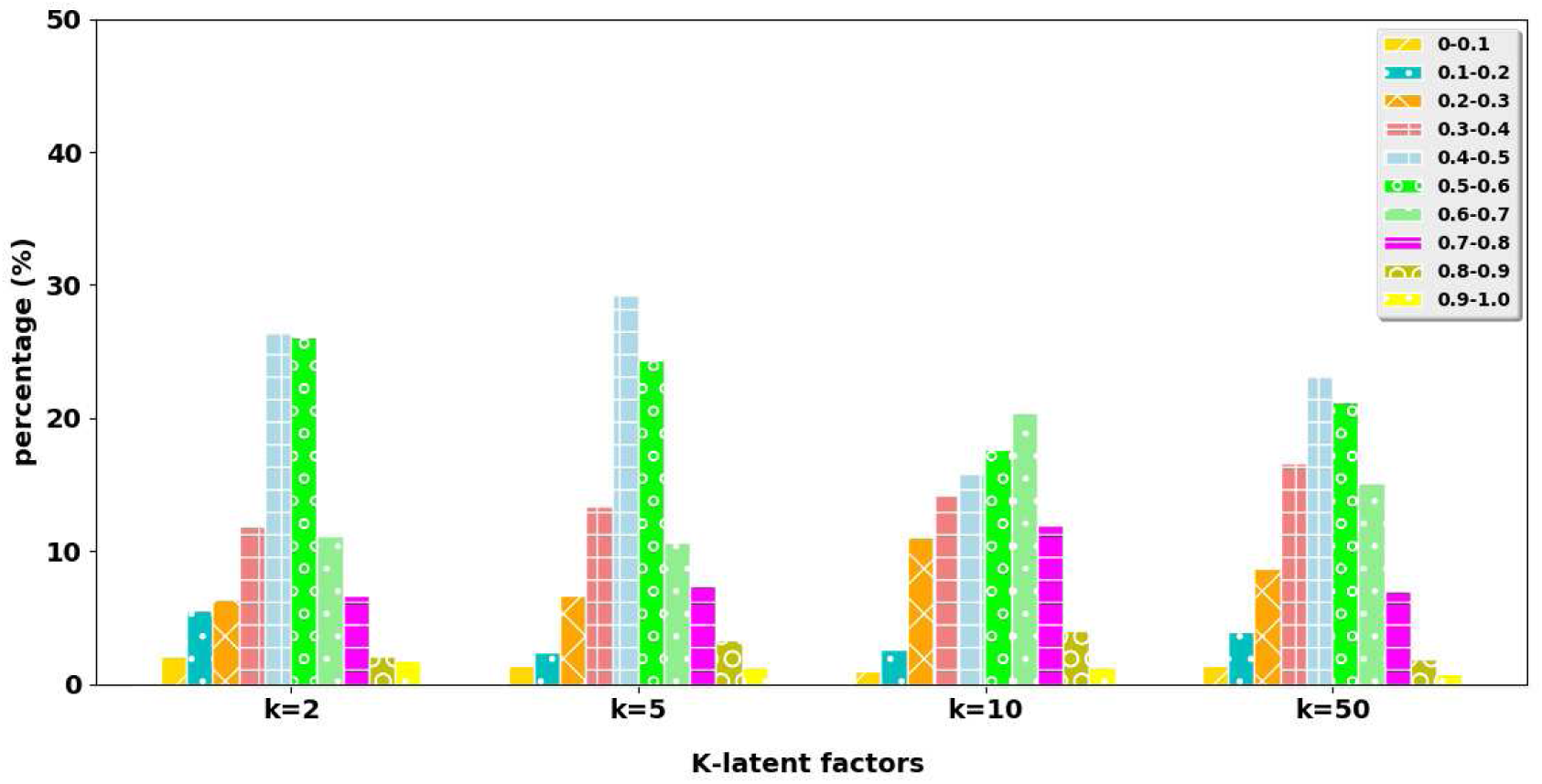
Experiment 3: distribution of values (in 10 bins after min-max scaling) of gene latent factor at four choices of *K*.

**Figure G.4:**
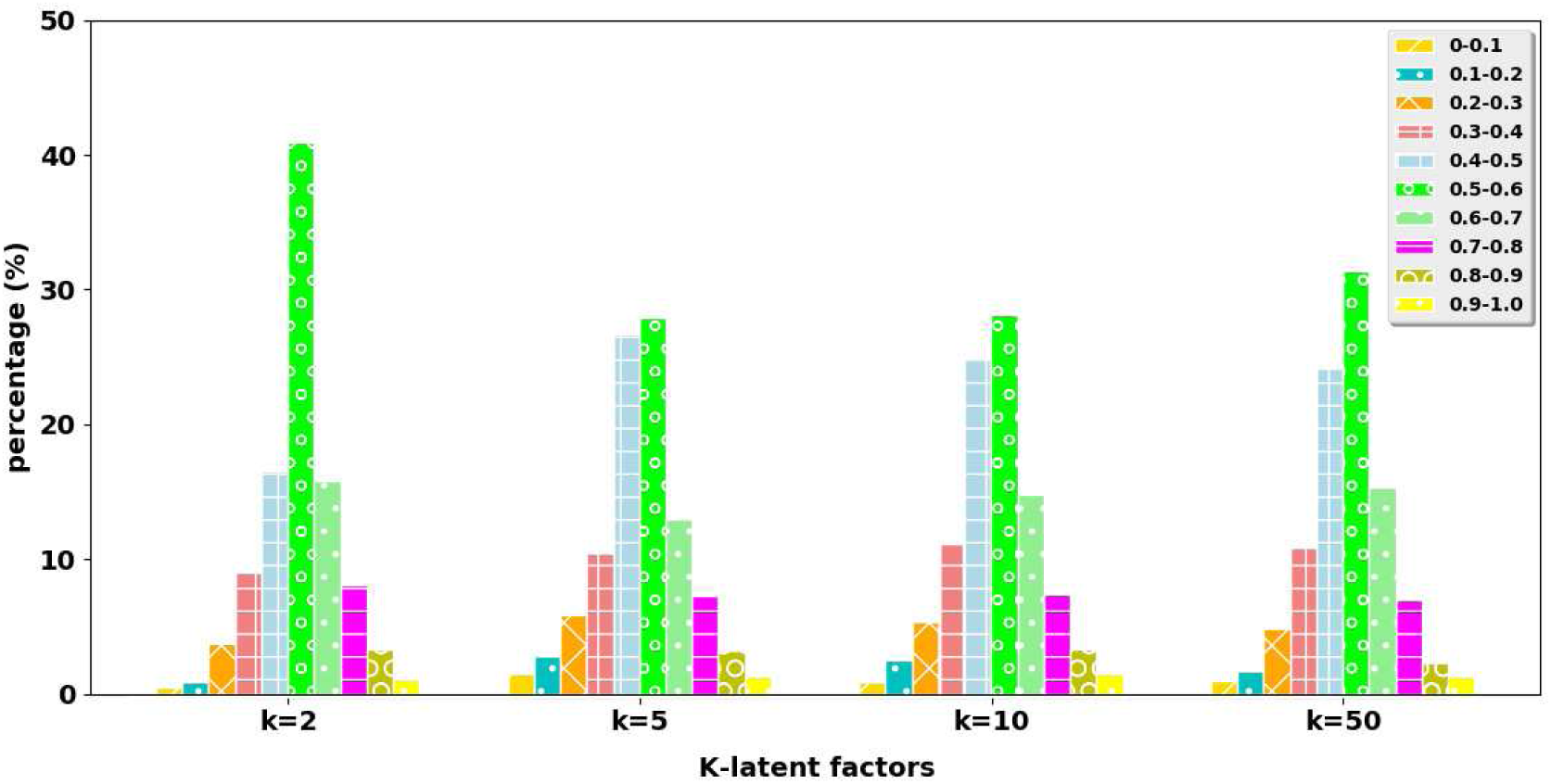
Experiment 4: distribution of values (in 10 bins after min-max scaling) of gene latent factor at four choices of *K*.

**Table G.1:** Experiment 1: distribution of values (in 10 bins after min-max scaling) of latent factors for all entities at four choices of *K*.

| Gene | 0-0.1 | 0.1-0.2 | 0.2-0.3 | 0.3-0.4 | 0.4-0.5 | 0.5-0.6 | 0.6-0.7 | 0.7-0.8 | 0.8-0.9 | 0.9-1 |
| --- | --- | --- | --- | --- | --- | --- | --- | --- | --- | --- |
| k=2 | 0.69% | 2.18% | 5.75% | 12.07% | 14.83% | 42.41% | 12.87% | 6.21% | 2.07% | 0.92% |
| k=5 | 1.43% | 3.72% | 7.36% | 17.70% | 39.68% | 15.22% | 8.55% | 4.14% | 1.52% | 0.69% |
| k=10 | 0.83% | 2.69% | 6.32% | 13.08% | 34.37% | 22.90% | 11.84% | 5.15% | 2.21% | 0.62% |
| k=50 | 0.85% | 2.24% | 5.83% | 11.84% | 26.12% | 29.65% | 13.41% | 6.50% | 2.61% | 0.97% |
| Hom del (PC) | 0-0.1 | 0.1-0.2 | 0.2-0.3 | 0.3-0.4 | 0.4-0.5 | 0.5-0.6 | 0.6-0.7 | 0.7-0.8 | 0.8-0.9 | 0.9-1 |
| k=2 | 8.33% | 5.00% | 11.67% | 16.67% | 11.67% | 11.67% | 8.33% | 16.67% | 5.00% | 5.00% |
| k=5 | 5.33% | 1.33% | 8.00% | 12.00% | 11.33% | 18.00% | 10.00% | 15.33% | 10.67% | 8.00% |
| k=10 | 6.00% | 6.00% | 5.67% | 10.67% | 15.00% | 12.33% | 14.00% | 9.67% | 9.33% | 11.33% |
| k=50 | 8.13% | 5.93% | 10.53% | 11.33% | 12.80% | 14.40% | 12.20% | 10.07% | 7.53% | 7.07% |
| Het del (PC) | 0-0.1 | 0.1-0.2 | 0.2-0.3 | 0.3-0.4 | 0.4-0.5 | 0.5-0.6 | 0.6-0.7 | 0.7-0.8 | 0.8-0.9 | 0.9-1 |
| k=2 | 3.33% | 1.67% | 11.67% | 16.67% | 18.33% | 13.33% | 25.00% | 5.00% | 0.00% | 5.00% |
| k=5 | 6.00% | 6.00% | 14.00% | 12.67% | 12.67% | 17.33% | 14.00% | 6.00% | 7.33% | 4.00% |
| k=10 | 6.00% | 6.67% | 10.67% | 14.00% | 14.33% | 16.00% | 14.67% | 6.67% | 5.00% | 6.00% |
| k=50 | 7.07% | 6.47% | 9.20% | 13.80% | 15.53% | 14.47% | 13.20% | 7.73% | 5.87% | 6.67% |
| Mix del (PC) | 0-0.1 | 0.1-0.2 | 0.2-0.3 | 0.3-0.4 | 0.4-0.5 | 0.5-0.6 | 0.6-0.7 | 0.7-0.8 | 0.8-0.9 | 0.9-1 |
| k=2 | 5.00% | 15.00% | 13.33% | 10.00% | 15.00% | 16.67% | 3.33% | 6.67% | 5.00% | 10.00% |
| k=5 | 8.00% | 6.00% | 8.00% | 12.67% | 13.33% | 14.00% | 11.33% | 12.67% | 7.33% | 6.67% |
| k=10 | 6.67% | 9.33% | 13.33% | 14.00% | 13.00% | 15.00% | 9.33% | 7.67% | 5.00% | 6.67% |
| k=50 | 7.27% | 6.20% | 10.67% | 11.53% | 17.07% | 14.13% | 12.67% | 8.67% | 4.40% | 7.40% |
| Co-UE del (PC) | 0-0.1 | 0.1-0.2 | 0.2-0.3 | 0.3-0.4 | 0.4-0.5 | 0.5-0.6 | 0.6-0.7 | 0.7-0.8 | 0.8-0.9 | 0.9-1 |
| k=2 | 10.00% | 10.00% | 13.33% | 10.00% | 16.67% | 15.00% | 8.33% | 0.00% | 6.67% | 10.00% |
| k=5 | 7.33% | 6.00% | 4.00% | 7.33% | 17.33% | 12.00% | 16.00% | 16.67% | 6.00% | 7.33% |
| k=10 | 5.67% | 4.00% | 8.00% | 14.33% | 16.00% | 18.67% | 13.00% | 7.33% | 6.67% | 6.33% |
| k=50 | 6.47% | 5.33% | 9.07% | 12.53% | 13.60% | 15.87% | 14.13% | 10.00% | 6.47% | 6.53% |
| Inv del (PC) | 0-0.1 | 0.1-0.2 | 0.2-0.3 | 0.3-0.4 | 0.4-0.5 | 0.5-0.6 | 0.6-0.7 | 0.7-0.8 | 0.8-0.9 | 0.9-1 |
| k=2 | 3.33% | 3.33% | 5.00% | 11.67% | 10.00% | 13.33% | 15.00% | 10.00% | 15.00% | 13.33% |
| k=5 | 6.67% | 4.00% | 11.33% | 17.33% | 17.33% | 12.67% | 7.33% | 11.33% | 3.33% | 8.67% |
| k=10 | 6.67% | 5.33% | 8.67% | 10.67% | 13.00% | 13.33% | 15.00% | 14.67% | 4.67% | 8.00% |
| k=50 | 6.13% | 5.47% | 9.73% | 12.33% | 13.60% | 14.07% | 14.93% | 9.87% | 7.00% | 6.87% |

**Table G.2:** Experiment 2: distribution of values (in 10 bins after min-max scaling) of latent factors for all entities at four choices of *K*.

| Gene | 0-0.1 | 0.1-0.2 | 0.2-0.3 | 0.3-0.4 | 0.4-0.5 | 0.5-0.6 | 0.6-0.7 | 0.7-0.8 | 0.8-0.9 | 0.9-1 |
| --- | --- | --- | --- | --- | --- | --- | --- | --- | --- | --- |
| k=2 | 1.51% | 3.31% | 6.33% | 14.76% | 34.49% | 17.02% | 11.14% | 7.68% | 2.71% | 1.05% |
| k=5 | 1.45% | 2.35% | 7.65% | 12.11% | 26.33% | 26.93% | 13.61% | 5.06% | 3.25% | 1.27% |
| k=10 | 1.05% | 2.35% | 6.05% | 10.33% | 27.77% | 24.79% | 15.75% | 6.90% | 3.80% | 1.20% |
| k=50 | 1.93% | 5.71% | 9.49% | 15.93% | 20.43% | 16.69% | 13.92% | 9.42% | 4.14% | 2.33% |
| Pairwise Co-expr (PC) | 0-0.1 | 0.1-0.2 | 0.2-0.3 | 0.3-0.4 | 0.4-0.5 | 0.5-0.6 | 0.6-0.7 | 0.7-0.8 | 0.8-0.9 | 0.9-1 |
| k=2 | 5.33% | 8.00% | 10.67% | 18.00% | 19.67% | 15.33% | 12.67% | 5.33% | 3.33% | 1.67% |
| k=5 | 1.73% | 2.80% | 7.47% | 16.80% | 18.80% | 19.87% | 16.93% | 7.87% | 5.47% | 2.27% |
| k=10 | 2.47% | 5.87% | 8.87% | 14.13% | 19.13% | 17.33% | 15.47% | 8.80% | 5.27% | 2.67% |
| k=50 | 2.15% | 4.87% | 9.81% | 15.43% | 17.53% | 18.17% | 15.16% | 9.96% | 4.69% | 2.23% |
| Gene Expr (Patient) | 0-0.1 | 0.1-0.2 | 0.2-0.3 | 0.3-0.4 | 0.4-0.5 | 0.5-0.6 | 0.6-0.7 | 0.7-0.8 | 0.8-0.9 | 0.9-1 |
| k=2 | 0.42% | 3.81% | 15.95% | 20.47% | 7.49% | 2.00% | 1.58% | 6.28% | 21.95% | 20.05% |
| k=5 | 11.55% | 23.55% | 18.57% | 4.97% | 1.34% | 2.25% | 10.21% | 9.41% | 6.33% | 11.81% |
| k=10 | 18.19% | 21.88% | 13.00% | 5.64% | 2.75% | 6.60% | 4.00% | 6.07% | 11.17% | 10.71% |
| k=50 | 4.69% | 9.65% | 15.73% | 12.86% | 9.02% | 12.58% | 11.03% | 11.75% | 9.61% | 3.07% |
| SCNA (PC) | 0-0.1 | 0.1-0.2 | 0.2-0.3 | 0.3-0.4 | 0.4-0.5 | 0.5-0.6 | 0.6-0.7 | 0.7-0.8 | 0.8-0.9 | 0.9-1 |
| k=2 | 0.67% | 0.00% | 2.00% | 3.67% | 31.67% | 30.00% | 25.33% | 5.00% | 0.67% | 1.00% |
| k=5 | 0.93% | 0.93% | 6.13% | 23.73% | 18.93% | 15.60% | 21.20% | 9.33% | 2.27% | 0.93% |
| k=10 | 2.20% | 11.07% | 8.67% | 9.27% | 15.60% | 17.87% | 17.53% | 6.33% | 9.40% | 2.07% |
| k=50 | 1.92% | 8.96% | 15.55% | 20.01% | 13.36% | 12.12% | 10.93% | 9.27% | 5.11% | 2.77% |

**Table G.3:** Experiment 3: distribution of values (in 10 bins after min-max scaling) of latent factors for all entities at four choices of *K*.

| Gene | 0-0.1 | 0.1-0.2 | 0.2-0.3 | 0.3-0.4 | 0.4-0.5 | 0.5-0.6 | 0.6-0.7 | 0.7-0.8 | 0.8-0.9 | 0.9-1 |
| --- | --- | --- | --- | --- | --- | --- | --- | --- | --- | --- |
| k=2 | 2.11% | 5.57% | 6.33% | 11.90% | 26.36% | 26.05% | 11.14% | 6.63% | 2.11% | 1.81% |
| k=5 | 1.17% | 2.83% | 7.17% | 12.67% | 18.50% | 23.83% | 13.67% | 11.83% | 4.67% | 3.67% |
| k=10 | 0.99% | 2.59% | 11.02% | 14.16% | 15.87% | 17.65% | 20.42% | 11.96% | 4.07% | 1.27% |
| k=50 | 1.38% | 3.97% | 8.74% | 16.63% | 23.28% | 21.22% | 15.08% | 6.95% | 1.92% | 0.83% |
| Essentiality (PC) | 0-0.1 | 0.1-0.2 | 0.2-0.3 | 0.3-0.4 | 0.4-0.5 | 0.5-0.6 | 0.6-0.7 | 0.7-0.8 | 0.8-0.9 | 0.9-1 |
| k=2 | 1.25% | 3.33% | 4.58% | 12.92% | 22.08% | 17.50% | 15.00% | 10.42% | 11.25% | 1.67% |
| k=5 | 1.73% | 2.80% | 7.47% | 16.80% | 18.80% | 19.87% | 16.93% | 7.87% | 5.47% | 2.27% |
| k=10 | 2.42% | 3.75% | 7.25% | 11.58% | 18.75% | 19.33% | 18.25% | 10.58% | 5.08% | 3.00% |
| k=50 | 2.90% | 5.35% | 10.23% | 14.73% | 19.53% | 18.90% | 14.12% | 8.28% | 3.67% | 2.28% |
| Phylo Score (Species) | 0-0.1 | 0.1-0.2 | 0.2-0.3 | 0.3-0.4 | 0.4-0.5 | 0.5-0.6 | 0.6-0.7 | 0.7-0.8 | 0.8-0.9 | 0.9-1 |
| k=2 | 46.88% | 37.44% | 12.05% | 2.37% | 0.84% | 0.28% | 0.05% | 0.00% | 0.00% | 0.09% |
| k=5 | 2.79% | 6.98% | 11.40% | 20.23% | 21.63% | 13.72% | 8.84% | 6.28% | 4.42% | 3.72% |
| k=10 | 5.81% | 10.93% | 12.21% | 12.56% | 12.79% | 12.33% | 13.49% | 10.35% | 6.16% | 3.37% |
| k=50 | 3.30% | 4.95% | 8.79% | 13.88% | 17.30% | 18.74% | 14.23% | 9.77% | 5.42% | 3.60% |
| Gene Expr (Patient) | 0-0.1 | 0.1-0.2 | 0.2-0.3 | 0.3-0.4 | 0.4-0.5 | 0.5-0.6 | 0.6-0.7 | 0.7-0.8 | 0.8-0.9 | 0.9-1 |
| k=2 | 0.42% | 3.81% | 15.95% | 20.47% | 7.49% | 2.00% | 1.58% | 6.28% | 21.95% | 20.05% |
| k=5 | 12.19% | 5.62% | 1.67% | 0.35% | 0.43% | 0.73% | 1.45% | 7.29% | 25.13% | 45.13% |
| k=10 | 22.11% | 14.73% | 8.41% | 3.37% | 1.20% | 0.75% | 1.16% | 5.13% | 17.74% | 25.40% |
| k=50 | 6.45% | 12.41% | 19.00% | 12.54% | 5.87% | 7.56% | 9.01% | 9.26% | 11.50% | 6.41% |
| SCNA (PC) | 0-0.1 | 0.1-0.2 | 0.2-0.3 | 0.3-0.4 | 0.4-0.5 | 0.5-0.6 | 0.6-0.7 | 0.7-0.8 | 0.8-0.9 | 0.9-1 |
| k=2 | 1.00% | 0.00% | 1.00% | 1.00% | 10.33% | 55.00% | 26.67% | 3.67% | 0.33% | 1.00% |
| k=5 | 2.27% | 2.40% | 15.20% | 20.93% | 14.27% | 8.13% | 13.07% | 14.13% | 7.20% | 2.40% |
| k=10 | 2.00% | 2.67% | 8.87% | 13.40% | 21.20% | 19.60% | 14.67% | 12.87% | 3.20% | 1.53% |
| k=50 | 3.31% | 8.59% | 9.59% | 11.09% | 16.31% | 18.52% | 16.31% | 11.69% | 3.41% | 1.19% |

**Table G.4:** Experiment 4: distribution of values (in 10 bins after min-max scaling) of latent factors for all entities at four choices of *K*.

| Gene | 0-0.1 | 0.1-0.2 | 0.2-0.3 | 0.3-0.4 | 0.4-0.5 | 0.5-0.6 | 0.6-0.7 | 0.7-0.8 | 0.8-0.9 | 0.9-1 |
| --- | --- | --- | --- | --- | --- | --- | --- | --- | --- | --- |
| k=2 | 0.45% | 0.90% | 3.77% | 9.04% | 16.57% | 40.96% | 15.81% | 8.13% | 3.31% | 1.05% |
| k=5 | 1.51% | 2.77% | 5.84% | 10.48% | 26.69% | 27.95% | 12.95% | 7.29% | 3.19% | 1.33% |
| k=10 | 5.00% | 1.33% | 10.00% | 13.00% | 27.00% | 5.33% | 4.67% | 10.00% | 17.33% | 6.33% |
| k=50 | 7.47% | 5.20% | 4.00% | 9.47% | 17.80% | 16.80% | 15.47% | 12.67% | 5.53% | 5.60% |
| Co-Expr (PC) | 0-0.1 | 0.1-0.2 | 0.2-0.3 | 0.3-0.4 | 0.4-0.5 | 0.5-0.6 | 0.6-0.7 | 0.7-0.8 | 0.8-0.9 | 0.9-1 |
| k=2 | 3.33% | 10.00% | 35.00% | 3.33% | 21.67% | 15.00% | 1.67% | 0.00% | 5.00% | 5.00% |
| k=5 | 22.67% | 3.33% | 1.33% | 4.00% | 29.33% | 22.67% | 8.67% | 0.00% | 2.67% | 5.33% |
| k=10 | 5.00% | 1.33% | 10.00% | 13.00% | 27.00% | 5.33% | 4.67% | 10.00% | 17.33% | 6.33% |
| k=50 | 2.90% | 5.35% | 10.23% | 14.73% | 19.53% | 18.90% | 14.12% | 8.28% | 3.67% | 2.28% |
| ME (PC) | 0-0.1 | 0.1-0.2 | 0.2-0.3 | 0.3-0.4 | 0.4-0.5 | 0.5-0.6 | 0.6-0.7 | 0.7-0.8 | 0.8-0.9 | 0.9-1 |
| k=2 | 50.00% | 0.00% | 0.00% | 0.00% | 0.00% | 0.00% | 0.00% | 0.00% | 0.00% | 50.00% |
| k=5 | 50.00% | 0.00% | 0.00% | 0.00% | 0.00% | 0.00% | 0.00% | 0.00% | 0.00% | 50.00% |
| k=10 | 50.00% | 0.00% | 0.00% | 0.00% | 0.00% | 0.00% | 0.00% | 0.00% | 0.00% | 50.00% |
| k=50 | 50.00% | 0.00% | 0.00% | 0.00% | 0.00% | 0.00% | 0.00% | 0.00% | 0.00% | 50.00% |
| Pathway (PC) | 0-0.1 | 0.1-0.2 | 0.2-0.3 | 0.3-0.4 | 0.4-0.5 | 0.5-0.6 | 0.6-0.7 | 0.7-0.8 | 0.8-0.9 | 0.9-1 |
| k=2 | 1.88% | 4.38% | 11.88% | 15.63% | 20.63% | 23.75% | 9.38% | 7.50% | 2.50% | 2.50% |
| k=5 | 3.25% | 4.25% | 10.50% | 18.75% | 21.50% | 15.25% | 10.50% | 8.50% | 4.50% | 3.00% |
| k=10 | 3.50% | 5.25% | 9.13% | 15.00% | 17.50% | 17.88% | 16.50% | 9.25% | 3.25% | 2.75% |
| k=50 | 3.13% | 5.30% | 9.70% | 14.73% | 17.68% | 17.48% | 14.13% | 9.20% | 4.78% | 3.90% |
| PPI (PC) | 0-0.1 | 0.1-0.2 | 0.2-0.3 | 0.3-0.4 | 0.4-0.5 | 0.5-0.6 | 0.6-0.7 | 0.7-0.8 | 0.8-0.9 | 0.9-1 |
| k=2 | 4.00% | 7.00% | 6.00% | 12.00% | 14.00% | 19.00% | 15.00% | 14.00% | 6.00% | 3.00% |
| k=5 | 4.40% | 6.80% | 12.40% | 14.40% | 12.00% | 15.60% | 17.60% | 9.60% | 3.20% | 4.00% |
| k=10 | 6.80% | 7.60% | 12.40% | 16.00% | 15.40% | 14.40% | 9.60% | 9.20% | 4.00% | 4.60% |
| k=50 | 5.32% | 5.96% | 10.72% | 15.08% | 14.76% | 15.60% | 13.56% | 9.04% | 5.96% | 4.00% |
| Prot-Complex (PC) | 0-0.1 | 0.1-0.2 | 0.2-0.3 | 0.3-0.4 | 0.4-0.5 | 0.5-0.6 | 0.6-0.7 | 0.7-0.8 | 0.8-0.9 | 0.9-1 |
| k=2 | 7.14% | 17.86% | 3.57% | 14.29% | 21.43% | 7.14% | 3.57% | 3.57% | 7.14% | 14.29% |
| k=5 | 14.29% | 7.14% | 8.57% | 10.00% | 7.14% | 8.57% | 17.14% | 11.43% | 2.86% | 12.86% |
| k=10 | 11.43% | 7.86% | 7.86% | 9.29% | 11.43% | 12.86% | 11.43% | 12.14% | 5.71% | 10.00% |
| k=50 | 11.71% | 5.57% | 11.29% | 12.00% | 11.86% | 11.86% | 9.29% | 9.14% | 5.71% | 11.57% |

## Footnotes

1 http://gdac.broadinstitute.org/

